# The Effect of Acute Pharmacological Inhibition of Urokinase Plasminogen Activator and Neuropsin Extracellular Proteases on Neuronal Networks *in vitro*

**DOI:** 10.1101/263616

**Authors:** Sebastiaan Van De Vijver, Stephan Missault, Jeroen Van Soom, Pieter Van der Veken, Koen Augustyns, Jurgen Joossens, Stefanie Dedeurwaerdere, Michele Giugliano

## Abstract

Neuronal networks are surrounded by the extracellular matrix (ECM), which functions both as a scaffold and as a regulator of neuronal function. The ECM is in turn dynamically altered through the action of serine proteases, which break down its constituents. This pathway has been implicated in the regulation of synaptic plasticity and of intrinsic excitability. Here, we determined the effects of acutely inhibiting two important regulators of the ECM, Urokinase Plasminogen Activator and Neuropsin, selectively and potently with the inhibitor UAMC-01162. Spontaneous electrophysiological activity was recorded from in vitro primary rat cortical cultures using microelectrode arrays. While inhibition at a low dosage had no significant effect, at elevated concentrations network bursting dynamics and functional connectivity were drastically altered. These results indicate that the serine protease inhibition affects neuronal and synaptic properties, likely through their actions on the ECM. We propose that in the acute phase, a transient increase of excitatory synaptic efficacy is compensated for by a downregulation of single-cell excitability.

## Introduction

The extracellular matrix (ECM) provides physical support and a stable environment to neurons *in vivo.* As a physical barrier, ECM may limit or refine structural connectivity, while at the same time also functioning as a critical regulator of synaptic plasticity (for a review, see [1]). The enzymatic removal of the ECM in *in vitro* cultures has been reported to facilitate the rearrangement of the neuronal connectivity profile [2]. Proteases are known to play an essential role in this regulation. Through their proteolytic action, they mould the structure of the ECM, which allows morphological changes to occur. Moreover, the extracellular proteolysis releases signalling molecules from the ECM, such as trans-synaptic proteins and growth factors, which are known to affect synaptic plasticity [3]. This evidence is especially strong for several serine proteases and matrix metalloproteinases, as reviewed in [3,4].

An important regulator of extracellular metabolism is the plasminogen-plasmin system. It is thought to be involved in structural remodelling and could thus affect neuronal connectivity. The urokinase-type plasminogen activator (uPA), a serine protease, converts plasminogen to plasmin, which in turn is responsible for the degradation of several extracellular proteins, both directly and indirectly. However, relatively little is known about the links between uPA and synaptic connectivity under physiological conditions in neuronal microcircuits. uPA has for instance been implicated in plasticity occurring during peripheral nerve regeneration [5–7], as well as in dendritic spine recovery following ischemic stroke [8]. In addition, uPA overexpression in transgenic animals has been associated with negative effects on learning and memory [9].

Neuropsin, also known as kallikrein-related peptidase 8 (KLK8), is another important serine protease that is expressed in an activity-dependent manner and secreted in the extracellular space as an inactive zymogen. Neuropsin is converted to its active form by NMDA-receptor-activation [10]. Through cleavage of target proteins, it activates a signalling pathway involving AMPA receptors, neuregulin-1, ErbB4, fibronectin, vitronectin, L1, integrin and L-type voltage-dependent Calcium ion channels [10–14]. Many of these targets are known for their clear involvement in structural and functional synaptic plasticity, and Neuropsin has been implicated by several lines of evidence in the regulation of synaptic plasticity [1]. Current data demonstrate that it modulates the early phase of long-term potentiation (LTP) [14,15], occurring independently of protein synthesis, that it enhances neurite outgrowth and fasciculation during development [16], that it primes synapses through tagging, making them more susceptible to persistent LTP [11,17], and that it strengthens GABAergic transmission [12]. Recently, Neuropsin has also been shown to affect the excitation-inhibition (E-I) balance in the hippocampus, through its modulation of parvalbumin-expressing interneuron activity via neuregulin-1 and ErbB4 signalling [13]. Moreover, an increased hyperexcitability to seizure-evoking stimuli [18] was reported in Neuropsin-deficient mice, with an associated altered E-I balance with decreased GABAergic interneuron activity and increased pyramidal neuron activity following administration of the convulsant kainic acid [13].

In this study, we investigated whether a highly selective and potent Neuropsin and uPA inhibitor, UAMC-01162, is capable of inducing changes in the network connectivity *in vitro* through its action on the ECM constituents. Such an effect is inferred from changes in the pattern of spontaneous electrophysiological activity, measured non-invasively and over extended periods of time from many neurons simultaneously, by means of substrate-integrated microelectrode arrays (MEAs; Fig. 1). This inhibitor consists of a diphenyl phosphonate group, which binds covalently to the serine alcohol in the catalytic centre of serine proteases and it results in a phosphorylated and inhibited enzyme, and a benzylguanidine group, which provides high selectivity and potency towards uPA and Neuropsin [19]. By measuring the electrical activity of neuronal networks by MEAs (Fig. 1B), underlying changes in connectivity can be assessed, while still preserving a functioning ECM [2,20,21]. Here, we present our experimental findings, referred to as the episodic and spontaneous synchronisation of “bursts” of action potentials (Fig. 1C-D) fired by neurons, and we demonstrate how the inhibition of serine proteases results in an altered network electrophysiology and effective connectivity.

**Figure 1.**
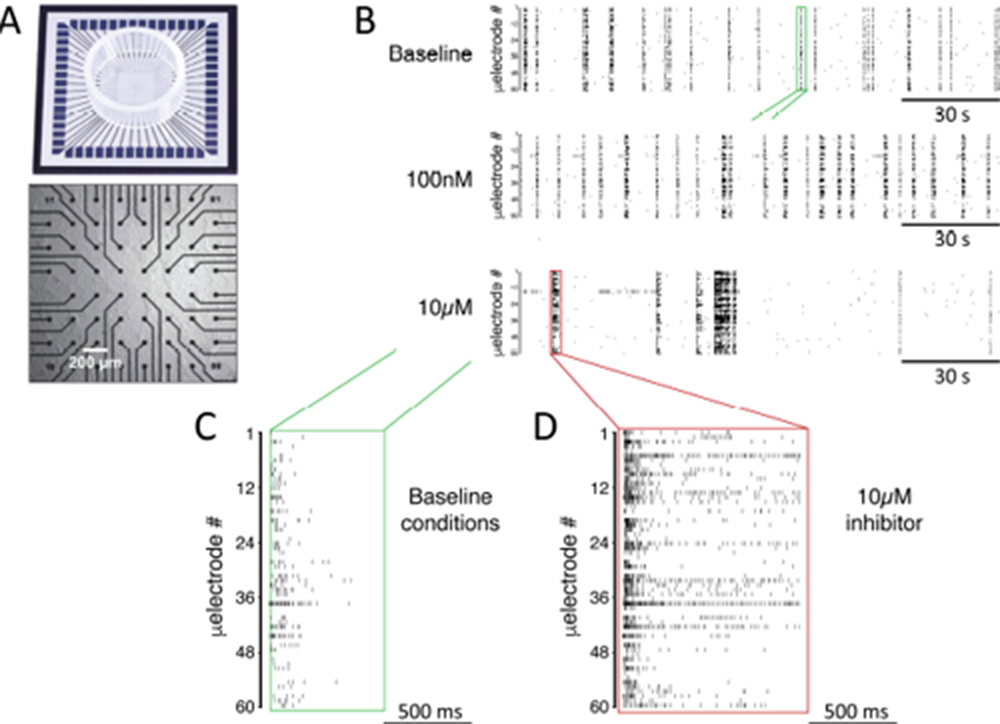
Experimental setup and a representative case of network-wide activity under inhibition of uPA/Neuropsin. (A) *In vitro* rat cortical networks were grown on substrate-integrated arrays of microelectrodes (MEAs) and their spontaneous electrical activity was monitored from up to 60 spatial locations. (B) A representative experiment is shown as a *raster plot:* each row refers to a distinct microelectrode, and each small marker represents the time of occurrence of a spike detected at that microelectrode. Three sample raster plots are shown from the same MEA, documenting baseline activity (top), low dose (middle), and high dose (bottom) of the compound tested. At high dose, the occurrence frequency of “bursts” of synchronous spikes is decreased, their timing variability increased, and their duration (compare C and D) increased. (C-D) A magnification over the bursts in B highlighted in green and red, reveal clearly the observed increase in burst duration.

## Materials and Methods

### Neuronal cultures

Primary cultures of mammalian neurons, dissociated from the postnatal rat neocortex, were prepared as described previously [22], in accordance with international and institutional guidelines on animal welfare. All procedures were approved by the Ethical Committee of the University of Antwerp (permission no. 2011_87) and licensed by the Belgian Animal, Plant and Food Directorate-General of the Federal Department of Public Health, Safety of the Food Chain and the Environment (license no. LA1100469). MEAs with a regular 8×8 arrangement of 60 titanium nitrate (TiN) microelectrodes, each with 30 μm diameter and 200 µm spacing (60MEA200/30iR-ITO-gr, MultiChannel Systems, Reutlingen, Germany), were employed in this study (Fig. 1A). Prior to cell seeding, each MEA was coated overnight with polyethyleneimine (0.1% wt/vol in milli-Q water at room temperature), and then extensively washed with milli-Q water and air-dried. After cell seeding, MEAs were maintained in a conventional incubator at 37 °C, 5% CO2, and 100% relative humidity (R.H.) (5215, Shellab, Cornelius, OR, USA). MEAs were sealed with fluorinated Teflon membranes (Ala-MEA-Mem, Ala Science, Farmingdale, NY, USA) during the entire duration of the experiments, to reduce the risk of contamination, prevent water evaporation and alteration of osmolarity, and ensure gas exchanges. After 8 days *in vitro* (DIV8), fresh and pre-warmed medium was added to reach a 1 ml final volume. From DIV10 onwards, half of the culture medium volume was replaced every 2 days with fresh, pre-warmed medium. All reagents were obtained from Sigma-Aldrich (St. Louis, MO, USA) or Life Technologies (Ghent, Belgium).

### Pharmacology and experimental protocol

The uPA/Neuropsin inhibitor (UAMC-01162) [19], dissolved in 2% DMSO, was added to the same cultures in two incremental “concentration steps”. One hour of baseline spontaneous recording activity was first acquired prior to applying the chemical compound. Subsequently, a small fraction of the culture medium was replaced with the uPA inhibitor solution, resulting in a final concentration of 100 nM, and the spontaneous electrical activity was recorded for one hour. Lastly, the final concentration of the compound was raised to 10 μM and the spontaneous electrical activity was recorded for one hour. At each step of compound addition to the bath, cultures were allowed to equilibrate for 10 minutes before starting each recording. Control cultures were treated under the same conditions, employing a DMSO solution without the inhibitor. The final DMSO concentration in the culture was 0.001% and 0.1% for the two concentration steps, respectively.

### Electrophysiological Recordings

Between DIV27 and 30, extracellular electrical recordings were made using a commercial amplifier (MEA-1060-Up-BC, MultiChannel Systems GmbH, Reutlingen, Germany) with a 1–3000 Hz bandwidth and an amplification factor of 1200. The raw voltage waveforms acquired from each of the 60 microelectrodes were sampled at 25 kHz/channel and digitised at 16 bits by an A/D electronic board (MCCard, MultiChannel Systems), and stored on disk by the MCRack software (Multi Channel Systems) for subsequent analysis.

### Data Analysis and Statistics

All data processing was performed using custom-written MATLAB scripts (The MathWorks, Natik, MA, USA). Action potential and network-burst detection was carried out using the QSpike Tools package [23]. In short, spike detection by peak detection was preceded by a band-pass filtering of the raw traces, and the threshold was adapted on the basis of the median of the raw voltage amplitude. All subsequent analyses were performed using the resulting timing of each threshold-crossing event, representing the occurrence of a putative action potential at any given microelectrode of the array (Fig. 1B-D). The microelectrodes were considered “active” if they showed events with at least an occurrence frequency of 0.02 Hz. A “burst” was defined as a major event (Fig. 1C-D) containing spiking activity in at least 15% of active electrodes in a 5 ms time window, with an artificial refractory period for their detection of 50 ms. The burst on- and offsets were defined as the points in time around the burst peak activity, where the Gaussian-smoothed network-wide spike-time histogram (STH, estimated using bins of 1 ms) reached zero. Basic statistics such as burst frequency, duration and inter-burst intervals were derived from these detected events if at least 20 bursts were detected per recording. The time course of the firing rate during each burst (i.e. STH) was further examined in terms of its slope at onset and of spectral (dominant) frequency content following the burst peak, if present. In detail, the rising phase of the STH, prior to the peak in activity associated to each burst, was fit by a double exponential function with equation f(t) = a_0_ e^(b_0_-1)t^ + a_1_ e^(b_1_-1)t^ + d using the non-linear least-square optimisation method. The smallest of the two exponents resulting from the fit, b0 or b1, was chosen to describe the rate of increase of network firing during the recruitment phase of each network burst. This value was obtained from the average burst-profile, for each specific condition and MEA.

Oscillatory frequency content in the offset phase of each burst was apparent in some, but not all MEAs [24]. This was revealed by pre-processing the data and individually aligning each burst prior to averaging, maximising similarity, as described previously. Frequency-domain analysis was then performed to extract the “dominant” frequency of the oscillation, by estimating the spectrogram [25] of each STH. In detail, the Fast Fourier Transform was employed to extract the time-varying spectrum of frequencies contained in the STH [26], averaging over all bursts. The “dominant” component of the power spectrum was defined as the Fourier frequency corresponding to the highest peak in the spectrum, which exceeded the median value of the spectrum by at least three fold.

All the parameters extracted from the data were evaluated over 30 min time windows, in order to assess whether or not stationarity of the recording was affected by the chemical compound application. Indeed, the compound under study here could have acute, persistent or delayed effects and care was taken to investigate the time course of each parameter. Furthermore, given the inherent variability of neuronal cultures, the relative changes in each parameter with respect to baseline, and not the absolute values, were taken to be the most important observables. Thus, each parameter was normalised to its respective value during the baseline recording condition.

Effective connectivity was extracted from each experiment by means of conventional cross correlation analysis of spike times [27]. Briefly, for each spike detected at electrode A, the statistics of relative time delays of all the spikes detected at electrode B were computed. This analysis was repeated over all spikes detected at A and restricted to delays smaller than 150 ms. Then the overall distribution of delays was estimated by a histogram with 3 ms bins and its profile was subsequently referred to as the cross correlogram, which quantified the effective coupling between each pair of MEA microelectrodes. Microelectrodes detecting too few spikes (n < 150) were excluded from this analysis.

As the persistent effects of the uPA/Neuropsin inhibition was of interest in this study, the cross-correlation analysis was limited to the last half hour of each recording session. For the baseline condition, the entire duration of the recordings was employed to increase statistical confidence. Each cross correlogram was normalised by the factor 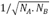, where N_A_ and N_B_ are the number of detected spikes for electrode A and B respectively. To represent the strength of the functional connection between each pair of microelectrodes, the peak value of the cross correlogram, C, was derived. Finally, the change in functional connectivity in response to the chemical inhibitor treatment was expressed as a ratio ∆C = C_after_/C_before_, where Cbefore is obtained from baseline conditions.

Statistical analysis was performed using MATLAB and it included the use of the Mann-Whitney U test to assess significant differences between the treated and control groups.

## Results

A hallmark feature of dissociated neuronal cultures is their tendency to spontaneously synchronise neuronal discharges in an episodic occurrence of “bursts” of electrical activity. Each burst lasts several hundreds of milliseconds [28] and recruits a large part of the network, as detected across many microelectrodes of the same MEA. During the onset of the burst, there is an explosive increase of the overall firing rate, as a dynamical reflection of the considerable structural recurrent excitation that effectively forms a positive feedback loop. The impact of the recruitment of inhibitory neurons is apparent at a later stage, together with the activation of several intrinsic and synaptic adaptation mechanisms, when the burst is suppressed and the network is ultimately silenced by an effective negative feedback loop [29]. The main features of these bursts, such as their duration and occurrence frequency, are therefore directly related to the mutual interaction and balance between excitation and inhibition in the network. In turn, the interactions between the excitatory and inhibitory subpopulations are directly determined by their synaptic connectivity and by their short- and long-term plasticity.

In our experiments, we investigated the effects of the selective and potent inhibition of uPA and Neuropsin, both key regulators of the ECM, on neuronal network activity *in vitro.* We repeated our experiments over n =10 distinct neuronal cultures where the uPA/Neuropsin inhibitor was bath applied, and we compared the results from several electrophysiological observables in n = 7 more neuronal cultures, employed as the control condition.

As a preliminary indication of the potential toxicity of our chemical compound, we examined the number of active microelectrodes, defined as those recording channels detecting neuronal activity above the minimal rate of 0.02 events/s (Fig. 2A). We found no significant effect caused by the uPA/Neuropsin inhibitor and we observed that all MEAs had an absolute number of active electrodes exceeding 90% (99.8 ± 1.8 *%* in control and 97.8 ± 1.1 *%* in low-dose treated MEAs, p = 0.909; 99.0 ± 3.2 % for control and 97.6 ± 1.5 % in high-dose treated MEAs, p = 0.762). These findings suggest that the compound does not markedly harm the neurons or negatively interfere with intrinsic membrane properties underlying synaptic input integration and the generation of action potentials.

**Figure 2.**
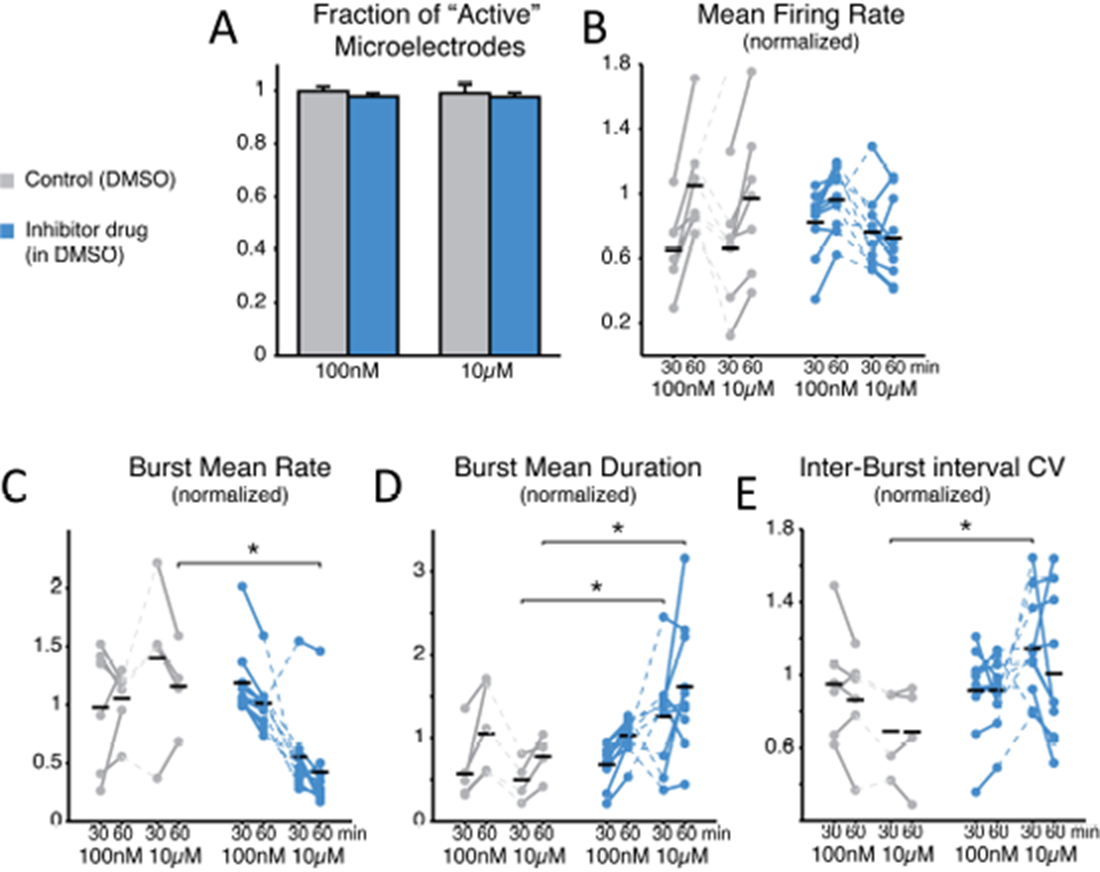
Inhibition of uPA/Neuropsin leads to a reduced burst occurrence, higher variability of inter-burst time intervals, and increased bursts duration. The compound does not affect neural viability as the average number of *active* microelectrodes (A), normalised to baseline condition, is left unaltered by the compound (blue: treated MEAs, n = 10; grey: control MEAs, n = 7; error bars are the s.e.m.). The mean firing rate over the entire MEA is displayed in (B), normalised to and estimated during the first and last 30 min time intervals of bath-application of the uPA/Neuropsin inhibitor, shows no significant difference (black horizontal bars are the group averages). Instead, the inhibitor alters the episodic occurrence of “bursts” of spikes, in cultures that display such a spontaneous activity pattern (10 treated and 6 controls at 100 nM, and 10 treated with 4 controls at 10 μ M): the burst mean rate, normalised to baseline, decreases particularly at the high dose of the inhibitor (C). Correspondingly, the burst duration increases significantly (D) at the high dose. Finally, the coefficient of variation of the interburst time interval distribution (E), normalised to baseline, increases particularly at the high dose of the inhibitor.

To gain an insight into the compound's effect on the structural synaptic connectivity, some basic observables were chosen to quantify neuronal firing and network bursting from the recorded spike trains under each condition (Fig. 2B-E). Despite the potency and strong selectivity of action, the uPA/Neuropsin inhibitor did not induce any appreciable effect when used at a low concentration (100nM). In fact, when expressing all the observables as normalised to baseline conditions, in the final 30 min period the firing rate was 1.049 ± 0.124 in control, 0.962 ± 0.058 for the treated group (p = 0.0962); the burst rate was 1.055 ± 0.110 in control, 1.013 ± 0.074 in treated MEAs (p = 0.313); the burst duration was 1.049 ± 0.217 in control, 1.028 ± 0.068 in treated cultures (p = 0.958); and the coefficient of variation of the inter-burst intervals was 0.853 ± 0.113 in control, 0.916 ± 0.061 in treated cultures (p = 0.875). However, when used at a high final concentration (10 μM), the periodic character of the *in vitro* network bursting changed significantly (Fig. 2C-E). This was the case, despite that neurons individually had similar firing rates (Fig. 2B; 0.661 ± 0.135 in controls and 0.762 ± 0.073 in treated cultures during the first 30 min after treatment, p = 0.601; 0.970 ± 0.177 in controls and 0.725 ± 0.081 in treated cultures in the second 30 min after treatment, p = 0.315). In fact, the occurrence frequency of the bursts decreased (Fig. 2C; 1.160 ± 0.187 in controls and 0.422 ± 0.120 in treated cultures, p = 0.014 < 0.05), while their duration increased (Fig. 2D; in the first 30 min, 0.498 ± 0.130 in controls and 1.259 ± 0.187 in treated cultures, p = 0.036 < 0.05; in the second 30 min, 0.778 ± 0.133 in controls and 1.619 ± 0.243 in treated cultures, p = 0.024 < 0.05), and their timing became more irregular, as indicated by an increase in the coefficient of variation of the inter-burst interval (Fig. 2E; 0.690 ± 0.119 in controls and 1.145 ± 0.090 in the treated group, p = 0.024 < 0.05).

Note that the control cultures displayed proportionally a qualitatively larger inter-individual variation in the firing rate and the burst frequency, compared to their baseline activity. The decreased burst frequency and increased irregularity of burst timings was significant (p < 0.05) for the second and first 30 min after drug administration, respectively.

When examined in its time course, the recruitment rate of the neuronal network during the burst initiation followed an exponential growth (Fig. 3A) [30]. When quantified by best fit of a double exponential functional, the recruitment rate (Fig. 3A, red trace) of both control and treated cultures showed similar rise times (Fig. 3B; during the final 30 min period, 0.987 ± 0.012 in controls and 0.995 ± 0.007 in cultures with low dose, p = 0.962, and with high dose controls 0.976 ± 0.016 and 1.015 ± 0.013 for the treated group, p = 0.070). As soon as successive bursts, recorded in the same MEA, were aligned for maximal similarity of their temporal profile and averaged (see the Methods) [24], a prominent oscillation in instantaneous rate of neuronal discharge became apparent during the falling phase of the bursts (e.g. Fig. 3A) in several but not all cultures. This is apparent from the structure of the power spectrum (Fig. 3C) and it is thought to arise from the interplay between the excitatory and inhibitory neuronal populations as in the Pyramidal-InterNeuron Gamma-range brain rhythms emergence *in vivo* (PING) [31]. When the “dominating” peaks (see the Methods) of the power spectrum were extracted (Fig. 3C), a significant increase in their Fourier frequency location could be observed, but only half an hour after the application of the inhibitor at the high dose (Fig. 3D; 0.824 ± 0.053 in controls and 1.701 ± 0.428 in the treated group, p = 0.028 < 0.05). As not all cultures revealed a clear-cut emergence of a dominant oscillation in the discharge rate of neurons during the bursts, cultures lacking oscillations in their corresponding baseline recordings were excluded from the analysis.

**Figure 3.**
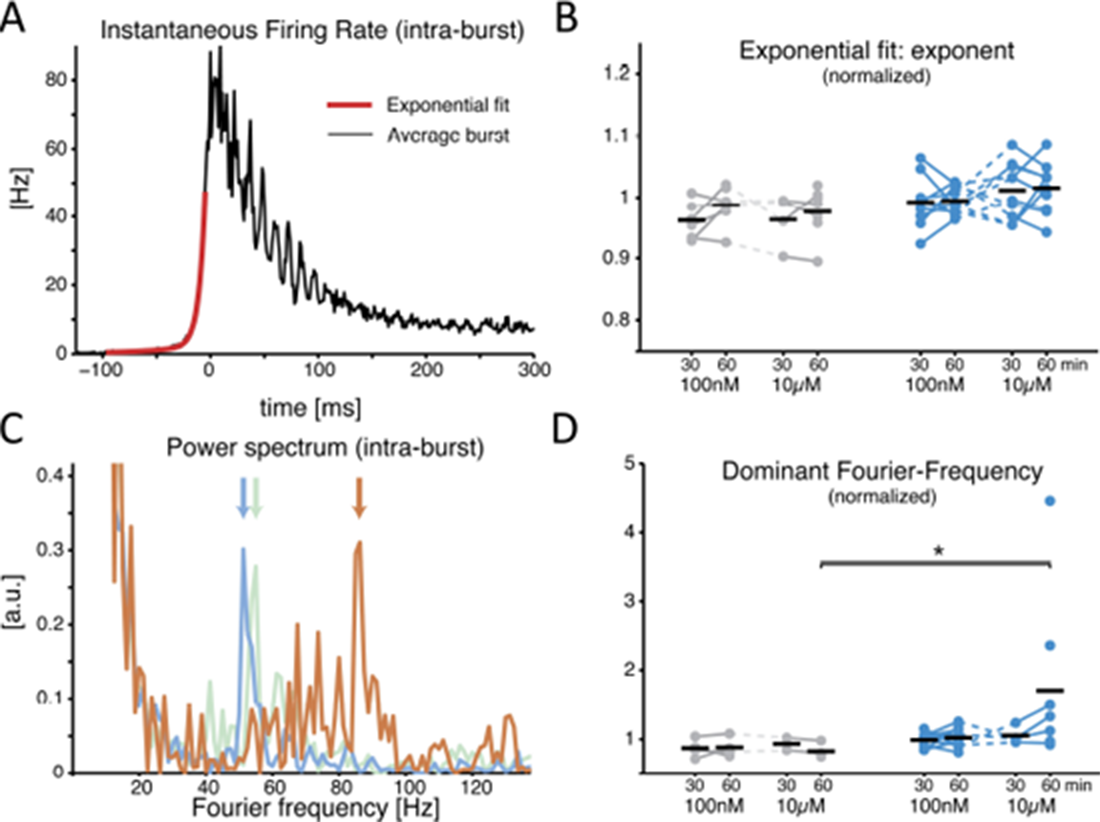
The uPA/Neuropsin inhibitor alters the features of the network burst at its offset, not onset. A representative example of the average instantaneous firing rate during a burst is displayed (A), estimated by a spike time histogram (STH). At onset the time course is best fit by an exponential function, while at offset it is often (not always) characterised by intraburst oscillations. When the onset is quantified, (B) by comparing the exponent of the best fit rising phase of the STH, normalised to baseline, the application of the inhibitor had no effect (black horizontal bars indicate the group averages). The power spectrum of the offset of the STH (C) quantifies an obvious transient oscillatory behaviour: coloured traces show the power spectrum density (PSD) for the baseline (grey), 100 nM (blue), and 10 μ M (orange) for the same MEA, and the location by a coloured arrow of the *dominant frequency.* When the latter is normalised to baseline and compared across conditions, a significant increase is found at the high dose of the inhibitor (D).

Finally, to quantify the effects of uPA/Neuropsin inhibition on synaptic connectivity, the strength of functional connections (see the Methods) was estimated in both the culture groups upon correlation analysis of the spike trains detected at distinct microelectrodes within the same MEAs, as in [32]. As displayed for a representative experiment (Fig. 4A), functional connectivity was significantly altered by the pharmacological inhibition of uPA/Neuropsin when compared to the (physiological) spontaneous redistribution of the coupling strength over the same amount of time [32]. When quantified over the entire set of experiments, the change in coupling strength (ΔC) was referred to baseline conditions and averaged across all possible microelectrode pairs (Fig. 4B). A highly significant increase in ∆C was observed in its mean (Fig. 4C; -0.214 ± 0.064 in controls and 0.063 ± 0.041 in the treated group, p = 0.000103 < 0.001) although not in its dispersion (Fig. 4D; 0.127 ± 0.015 in controls and 0.125 ± 0.010 in treated cultures, p = 0.669), suggesting that stronger effective coupling appeared over the course of the time interval following the application of the inhibitor.

**Figure 4.**
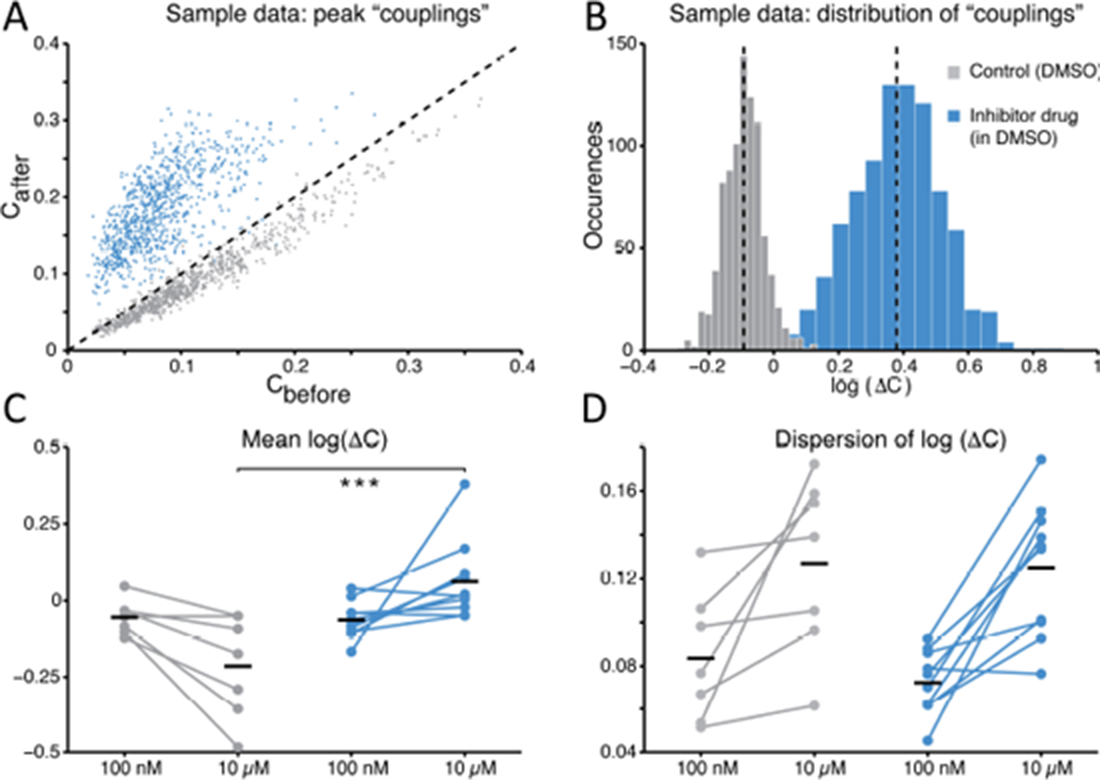
The average *functional coupling* across any microelectrode pair appears strengthened under uPA/Neuropsin inhibition. When compared to control conditions (grey), the strength of *functional coupling* increases (A) with the inhibitor at the high dose (blue). Each dot in the scatter plot corresponds to one of the possible microelectrode pairs and displays the value of the peak of the spike cross-correlogram: dots above the unitary slope line imply an increase in strength with respect to baseline. The actual distribution of values, in the same representative example of panel (A), can be better displayed in logarithmic scale (base 10) while expressing the change (B) after the dose referred to the baseline ΔC = C_after_/C_before_): the black dotted vertical lines are the mean values of the distributions (control *versus* high dose) and are significantly different. Across all our experiments (C), 10 μ M of the uPA/Neuropsin inhibitor significantly strengthened the functional couplings between microelectrodes compared to the control, while the latter tends to even decrease over time (black horizontal bars indicate the group averages). The standard deviation or *dispersion* (D) of the log10 ΔC) distributions increases over time, but in the same manner comparing the effect of the compound and the control.

## Discussion

The ECM is a key player in the regulation of intrinsic excitability and structural synaptic connectivity, although the involvement of its different molecular constituents is largely unknown. Proteolytic regulation of ion channels is associated with the ECM [33] and the mechanisms underlying action potential generation (e.g. voltage-gated sodium channels) are also implicated in adhesion, migration, path finding, and transcription [34].

We investigated the indirect effect of a selective and potent serine protease inhibitor on network-level electrophysiology, targeting two important regulators of the ECM: uPA and Neuropsin. We observed significant alterations in the collective emergent activity *in vitro* within just 1h following the application of the inhibitor, consistent with the rapid *in vivo* action of ECM proteases (i.e. over minutes to hours [33]). Our observations seem to point to both synaptic connectivity and to intrinsic single-cell excitability, in line with our current understanding of ECM proteases (see [33] for a review).

The increase of the average burst duration, the decrease in the burst occurrence, the increase in inter-burst variability, and the increase in the oscillation frequency within bursts indicate a weakening of synaptic excitation and an increase in synaptic inhibition. A weakened excitation and a reduction in inhibition would allow the recurrent excitation to ignite a burst less frequently and more randomly over time and, once started, to let it carry on for longer while delaying its termination. This has been illustrated in an early study by Giugliano and collaborators [35] and also by Gross and collaborators, who boosted inhibition through administration of GABA and decreased burst duration [36]. A shift towards inhibition is also predicted by a mean-field model of the network activity, leading to faster intra-burst oscillations [37]. There, an increase of the efficacy of inhibitory connections or their numbers increases the intra-burst frequency oscillation.

However, this is in part contradicting the significant increase in the action potential firing synchrony through cross-correlation analysis, despite being only an indirect measure of effective and not anatomical coupling. An alternative hypothesis might be that the intrinsic membrane excitability was also reduced together with an increase of the excitatory coupling, so that the random episodic occurrence of a burst by means of recruitment of network target occurs less frequently but with higher synchrony.

The similar exponential behaviour of the early, i.e. rising, phase of the bursts suggests that the same avalanche-like mechanism of burst initiation by recruitment of neurons by recurrent excitatory connections is present. The termination of the bursts and its delayed occurrence, after the application of the inhibitor suggests that the same intrinsic mechanisms (e.g. calcium-dependent outward potassium currents) continued to operate but that the threshold for their intervention was reached later given the lower single-cell excitability.

The increased frequency of intra-burst oscillations in the instantaneous firing rate of the network is consistent with an increase in the excitatory synaptic efficacy as well as with an increase in the inhibitory synaptic efficacy [37]. Inhibition, when recruited by excitatory neurons, operates with a dampening effect on the network's explosive behaviour during a burst. Ultimately, the balance between excitation and inhibition in the network seems to have been altered. However, we note that a shift in the E-I balance has not always led to consistent changes in burst dynamics. For example, in 2007 Chiappalone and collaborators showed that both the duration of bursting and the interval between the bursts, which is inversely related to bursting frequency, increase in response to the GABA*_A_* antagonist bicuculline [38]. The resulting shift towards excitation produces the expected effect on burst duration. The measured activity as visualised in their raster plots is also remarkably similar to the activity observed in the present study. Additionally, a more recent study by Odawara and collaborators shows that the relationship between the E-I balance and burst frequency or burst duration changes over time [39]. At an earlier developmental age, their model system responds to bicuculline treatment through an increase in burst frequency, while the burst duration undergoes a small and non-significant increase. In contrast, later in development the same networks exposed to the same treatment only show a modest non-significant increase in burst frequency while burst duration is markedly longer. How a shift in the E-I balance affects particular aspects of the bursting dynamics thus seems to depend on the state of the network and these are unreliable for deducting the direction of E-I balance shift.

Evidence from the literature supports a similar shift in response to decreased uPA or Neuropsin levels. Complete deficiency of Neuropsin has been shown to shift the E-I balance *in vivo* towards excitation, in the hippocampus of mice treated with kainic acid, thus leading to a more severe progression of the status epilepticus than in control [13]. Similarly, in an animal model of epilepsy, a pathological condition associated with strong recurrent excitation, we have recently found that the frequency of seizures is higher in epileptic rats that show a more pronounced decrease in active uPA and Neuropsin [40]. While our results do not unanimously point to a shift of the E-I balance in either direction, the inhibition of uPA and Neuropsin certainly had a profound effect on the dynamics of the network.

Furthermore, at the lowest concentration (100 nM) the compound demonstrated no noticeable effects, despite being several times above the IC50 for uPA, Neuropsin (KLK8) and KLK4 (i.e. 21, 15, and 44 nM, respectively). It is likely that the compound is capable of inhibiting additional serine proteases. In human tissue, the inhibitor has also shown affinity in the nanomolar range for cathepsin G, tryptase, and matriptase. Additionally, uPA expression levels were shown to be low in murine CNS under normal conditions [41], only being upregulated during pathological conditions [42,43].

While in the long-term, the structural connectivity of the network might ultimately result in a diminished excitatory coupling and an increase in the number of inhibitory neurons as observed in ultrastructural studies of Neuropsin-deficient animals, in the short-term as observed in our study, several synaptic and intrinsic homeostatic mechanisms may operate with opposing behaviour: a downregulation of single-cell excitability to compensate for the transiently increased excitatory coupling.

We have demonstrated that manipulation of the ECM with the inhibitor changed the connectivity of the network, resulting in an altered balance between excitation and inhibition. Future studies are required to further identify the targets and pathways through which this inhibitor can change the pattern of activity in the network. A more detailed identification of all the molecular players in the ECM and of their roles in regulating neuronal connectivity is needed to further unravel the mechanisms of brain wiring.

## Acknowledgements

We are grateful to Mr D. Van Dyck, Mrs G. Van de Vijver, and Mr M. Wijnants for excellent technical assistance and helpful discussions. Financial support from the 7th FP of the European Commission (FP7-FP7-PEOPLE-IAPP grant n. 286403, FP7-ICT-FET grant n. 284801), the H2020 (H2020-PEOPLE-ETN grant n. 642881), the ERA-NET (NEURON II project “TBI Epilepsy”), the Interuniversity Attraction Poles Program (IUAP) of the Belgian Science Policy Office, and the University of Antwerp Research Fund (BOF) is kindly acknowledged. The funders had no role in study design, data collection and analysis, decision to publish, or preparation of the manuscript.

## References

1. Mukhina I, Korotchenko SA, Dityatev AE. Extracellular matrix molecules, their receptors, and extracellular proteases as synaptic plasticity modulators. Neurochem J. 2012;29(2): 106–17.

2. Bikbaev A, Frischknecht R, Heine M. Brain extracellular matrix retains connectivity in neuronal networks. Sci Rep. 2015;5(1): 14527.

3. Tsilibary E, Tzinia A, Radenovic L, Stamenkovic V, Lebitko T, Mucha M, et al. Neural ECM proteases in learning and synaptic plasticity. In: Dityatev A, Wehrle-Haller B, Pitkänen A, editors. Brain Extracellular Matrix in Health and Disease. 2014. p. 214: 135–57.

4. Shiosaka S. Serine proteases regulating synaptic plasticity. Anat Sci Int. 2004;79(3): 137–44.

5. Seeds NW, Mikesell S, Vest R, Bugge T, Schaller K, Minor KH. Plasminogen activator promotes recovery following spinal cord injury. Cell Mol Neurobiol. 2011;31(6):961–7.

6. Karagyaur M, Dyikanov D, Makarevich P, Semina E, Stambolsky D, Plekhanova O, et al. Non-viral transfer of BDNF and uPA stimulates peripheral nerve regeneration. Biomed Pharmacother. 2015;74:63–70.

7. Minor KH, Seeds NW. Plasminogen activator induction facilitates recovery of respiratory function following spinal cord injury. Mol Cell Neurosci. 2008;37:143–52.

8. Wu F, Catano M, Echeverry R, Torre E, Haile WB, An J, et al. Urokinase-type plasminogen activator promotes dendritic spine recovery and improves neurological outcome following ischemic stroke. J Neurosci. 2014;34(43):14219–32.

9. Meiri N, Masos T, Rosenblum K, Miskin R, Dudai Y. Overexpression of urokinase-type plasminogen activator in transgenic mice is correlated with impaired learning. Proc Natl Acad Sci U S A. 1994;91:3196–200.

10. Matsumoto-Miyai K, Ninomiya A, Yamasaki H, Tamura H, Nakamura Y, Shiosaka S. NMDA-dependent proteolysis of presynaptic adhesion molecule L1 in the hippocampus by neuropsin. J Neurosci. 2003;23(21):7727–36.

11. Ishikawa Y, Horii Y, Tamura H, Shiosaka S. Neuropsin (KLK8)-dependent and -independent synaptic tagging in the Schaffer-collateral pathway of mouse hippocampus. J Neurosci. 2008;28(4):843–9.

12. Tamura H, Kawata M, Hamaguchi S, Ishikawa Y, Shiosaka S. Processing of neuregulin-1 by neuropsin regulates GABAergic neuron to control neural plasticity of the mouse hippocampus. J Neurosci. 2012;32(37):12657–72.

13. Kawata M, Morikawa S, Shiosaka S, Tamura H. Ablation of neuropsin-neuregulin 1 signaling imbalances ErbB4 inhibitory networks and disrupts hippocampal gamma oscillation. Transl Psychiatry. 2017 Mar 7;7.

14. Tamura H, Ishikawa Y, Hino N, Maeda M, Yoshida S, Kaku S, et al. Neuropsin is essential for early processes of memory acquisition and Schaffer collateral long-term potentiation in adult mouse hippocampus in vivo. J Physiol. 2006;570(3):541–51.

15. Komai S, Matsuyama T, Matsumoto K, Kato K, Kobayashi M, Imamura K, et al. Neuropsin regulates an early phase of Schaffer-collateral long-term potentiation in the murine hippocampus. Eur J Neurosci. 2000;12:1479–86.

16. Oka T, Akisada M, Okabe A, Sakurai K, Shiosaka S, Kato K. Extracellular serine protease neuropsin (KLK8) modulates neurite outgrowth and fasciculation of mouse hippocampal neurons in culture. Neurosci Lett. 2002;321(3):141–4.

17. Ishikawa Y, Tamura H, Shiosaka S. Diversity of neuropsin (KLK8)-dependent synaptic associativity in the hippocampal pyramidal neuron. J Physiol. 2011;589(14):3559–73.

18. Davies B, Kearns IR, Ure J, Davies CH, Lathe R. Loss of hippocampal serine protease BSP1/neuropsin predisposes to global seizure activity. J Neurosci. 2001;21(18):6993–7000.

19. Joossens J, Ali OM, El-Sayed I, Surpateanu G, Der Van Veken P, Lambeir AM, et al. Small, potent, and selective diaryl phosphonate inhibitors for urokinase-type plasminogen activator with in vivo antimetastatic properties. J Med Chem. 2007;50:6638–46.

20. Dityatev A, Brückner G, Dityateva G, Grosche J, Kleene R, Schachner M. Activity-dependent formation and functions of chondroitin sulfate-rich extracellular matrix of perineuronal nets. Dev Neurobiol. 2006;67(5):570–88.

21. John N, Krügel H, Frischknecht R, Smalla KH, Schultz C, Kreutz MR, et al. Brevican-containing perineuronal nets of extracellular matrix in dissociated hippocampal primary cultures. Mol Cell Neurosci. 2006;31(4):774–84.

22. Reinartz S, Biro I, Gal A, Giugliano M, Marom S. Synaptic dynamics contribute to long-term single neuron response fluctuations. Front Neural Circuits. 2014;8(July).

23. Mahmud M, Pulizzi R, Vasilaki E, Giugliano M. QSpike tools: a generic framework for parallel batch preprocessing of extracellular neuronal signals recorded by substrate microelectrode arrays. Front Neuroinform. 2014;8(March).

24. Pulizzi R, Musumeci G, Van den Haute C, Van De Vijver S, Baekelandt V, Giugliano M. Brief wide-field photostimuli evoke and modulate oscillatory reverberating activity in cortical networks. Sci Rep. 2016;

25. Oppenheim A V, Schafer RW, Buck JR. Discrete Time Signal Processing. Book. 1999. p. 870.

26. Mattia M, Ferraina S, Del Giudice P. Dissociated multi-unit activity and local field potentials: A theory inspired analysis of a motor decision task. Neuroimage. 2010;52(3):812–23.

27. Knox CK. Detection of neuronal interactions using correlation analysis. Trends Neurosci. 1981;4(September) : 222–5.

28. Kamioka H, Maeda E, Jimbo Y, Robinson HPC, Kawana A. Spontaneous periodic synchronized bursting during formation of mature patterns of connections in cortical cultures. Neurosci Lett. 1996;206(2-3):109–12.

29. Robberechts Q, Wijnants M, Giugliano M, De Schutter E. Long-Term Depression at Parallel Fiber to Golgi Cell Synapses. J Neurophysiol. 2010;104(6):3413–23.

30. Eytan D, Marom S. Dynamics and effective topology underlying synchronization in networks of cortical neurons. J Neurosci. 2006;26(33):8465–76.

31. Wang X-J. Neurophysiological and Computational Principles of Cortical Rhythms in Cognition. Physiol Rev. 2010;90(3):1195–268.

32. Eytan D, Minerbi A, Ziv N, Marom S. Dopamine-induced dispersion of correlations between action potentials in networks of cortical neurons. J Neurophysiol. 2004;92(3): 1817–24.

33. Wójtowicz T, Brzdąk P, Mozrzymas JW. Diverse impact of acute and long-term extracellular proteolytic activity on plasticity of neuronal excitability. Front Cell Neurosci. 2015;9(313): 1–18.

34. Bourgognon J, Schiavon E, Salah-uddin H, Skrzypiec AE, Attwood BK, Shah RS, et al. Regulation of Neuronal Plasticity and Fear by a Dynamic Change in PAR1 – G Protein Coupling in the Amygdala. Mol Psychiatry. 2013; 18(10): 1136–45.

35. Giugliano M, Darbon P, Arsiero M, Lüscher H-R, Streit J. Single-Neuron Discharge Properties and Network Activity in Dissociated Cultures of Neocortex. J Neurophysiol. 2004;92(2):977–96.

36. Gross GW, Rhoades B, Jordan R. Neuronal networks for biochemical sensing. Sensors Actuators B. 1992;6:1–8.

37. Pulizzi R, Musumeci G, Van Den Haute C, Baekelandt V, Giugliano M. Wide-field photostimulation in in vitro cortical networks: prolonged selective activation of (CaMKIIα)-positive excitatory neurons and its impact on evoked reverberating responses. In 2013.

38. Chiappalone M, Vato A, Berdondini L, Koudelka-Hep M, Martinoia S. Network Dynamics and Synchronous Activity in Cultured Cortical Neurons. Int J Neural Syst. 2007;17(2):87–103.

39. Odawara A, Katoh H, Matsuda N, Suzuki I. Physiological maturation and drug responses of human induced pluripotent stem cell-derived cortical neuronal networks in long-term culture. Sci Rep. 2016;6:1–14.

40. Missault S, Peeters L, Amhaoul H, Thomae D, Van Eetveldt A, Favier B, et al. Decreased levels of active uPA and KLK8 assessed by [ 111 In]MICA-401 binding correlate with the seizure burden in an animal model of temporal lobe epilepsy. Epilepsia. 2017;58(9):1–11.

41. Sappino AP, Madani R, Huarte J, Belin D, Kiss JZ, Wohlwend A, et al. Extracellular proteolysis in the adult murine brain. J Clin Invest. 1993;92(2):679–85.

42. Lahtinen L, Lukasiuk K, Pitkänen A. Increased expression and activity of urokinase-type plasminogen activator during epileptogenesis. Eur J Neurosci. 2006;24(7):1935–45.

43. Gorter JA, Van Vliet EA, Rauwerda H, Breit T, Stad R, Van Schaik L, et al. Dynamic changes of proteases and protease inhibitors revealed by microarray analysis in CA3 and entorhinal cortex during epileptogenesis in the rat. Epilepsia. 2007;48(SUPPL. 5):53–64.

